# Mutations in the *ubiA* gene are the major mechanism of ethambutol resistance in *Mycobacterium avium*

**DOI:** 10.1101/2025.09.27.678926

**Authors:** Yuwei Qiu, Xu Dong, Dan Cao, Xiuzhi Jiang, Zhongkang Ji, Pusheng Xu, Yi Li, Kaijin Xu, Ying Zhang

**Affiliations:** State Key Laboratory for Diagnosis and Treatment of Infectious Diseases, National Clinical Research Center for Infectious Diseases, China-Singapore Belt and Road Joint Laboratory on Infection Research and Drug Development, National Medical Center for Infectious Diseases, Collaborative Innovation Center for Diagnosis and Treatment of Infectious Diseases, The First Affiliated Hospital, Zhejiang University School of Medicine; Yuhang Institute for Collaborative Innovation and Translational Research in Life Sciences and Technology

**Keywords:** *Mycobacterium avium*, ethambutol, antibiotic resistance, mutations, minimum inhibitory concentration

## Abstract

*Mycobacterium avium-intracellulare* complex (MAC) infections are the most common nontuberculous mycobacterial (NTM) infections, and ethambutol (EMB) is one of the main therapeutic agents used to treat MAC infections. However, the EMB resistance profile and its resistance mechanisms in *M. avium* remain poorly understood. This study determined the minimum inhibitory concentration (MIC) on 40 *M. avium* clinical strains, revealing that 97.5% (39/40) of the strains were intermediate (MIC=4 μg/mL) or resistant (MIC≥8 μg/mL) to EMB. One susceptible clinical isolate strain 245 (MIC=2 μg/mL) was picked for EMB-resistant mutant isolation, and a total of 121 resistant mutants were isolated and subjected to whole-genome sequencing or Sanger sequencing. Integrated analysis revealed that 94.21% (114/121) of the mutants carried mutations in the *ubiA* gene, which encodes DPPR (decaprenylphosphoryl-β-D-5-phosphoribose) synthase - an enzyme involved in cell wall biosynthesis that has been associated with high-level EMB resistance in *M. tuberculosis*. Complementation with the wild-type *ubiA* gene restored EMB susceptibility in EMB-resistant mutants and clinical strain 322 (EMB MIC=64 μg/mL) with *ubiA* mutation, reducing the MICs from 32 μg/mL to 4 μg/mL and from 64 μg/mL to 8 μg/mL, respectively. This study indicates that the *ubiA* mutation is the major mechanism of EMB resistance in *M. avium*, which should facilitate the development of molecular tests for rapid detection of resistance in this organism.

## INTRODUCTION

*Mycobacterium avium* complex (MAC) is a group of several slow-growing mycobacteria (SGM) and is commonly found in the environment such as water and soil (1, 2). MAC is primarily transmitted through aerosol inhalation, and can cause infections often in immunocompromised individuals (3). Patients who are aged, immunosuppressed, or with other respiratory diseases (such as cystic fibrosis, bronchiectasis and COPD) are at higher risk of developing severe clinical symptoms following MAC infections (4–6). MAC comprises various subspecies, including *M. avium*, *M. intracellulare*, *M. colombiense, M. timonense*, etc., with *M. avium* being one of the most prevalent species globally (7). Current treatment guidelines for MAC infections recommend a multi-drug combination therapy with macrolides as the core drug, along with ethambutol (EMB), rifampicin (RIF), aminoglycosides or other drugs (8).

EMB, as a key component in MAC treatment regimens, inhibits bacterial growth by interfering with arabinogalactan biosynthesis in the mycobacterial cell wall (9). Not only does its use significantly improve sputum conversion rates compared to RIF, streptomycin, or fluoroquinolones in the treatment of CLR-resistant MAC infections, but it also effectively prevents the development of macrolide resistance (10, 11). Although a previous study classified MAC strains as susceptible (MIC≤2 μg/mL), intermediate (MIC=4 μg/mL), or resistant (MIC≥8 μg/mL) to EMB, many studies indicated that the MICs of *M. avium* strains of EMB mainly are 8 μg/mL, while Clinical and Laboratory Standards Institute (CLSI) does not yet have EMB susceptibility or resistance breakpoints for *M. avium* (12–14). Furthermore, it has been suggested that when MAC isolates exhibit MICs≥8 μg/mL for both EMB and RIF, the clinical treatment success rate will decrease significantly, indicating the negative correlation between successful treatment outcome with resistance to EMB and RIF (15). Therefore, elucidating the mechanism of EMB resistance in MAC remains clinically significant. To date, *embB/embR aftA* and *ubiA* genes have been proved associated with EMB resistance in *M. tuberculosis* (MTB) (16–19), but the resistance mechanism in MAC is far less explored. Although previous studies have revealed that *embA*/*B* contribute to elevated EMB resistance in *M. avium* (20), clinical investigation indicated that *embB* mutations account for only about 4% of the EMB-resistant MAC clinical isolates (21), indicating other major EMB resistance mechanisms remain to be identified.

To better understand the mechanisms of EMB resistance in *M. avium*, here, we isolated EMB-resistant mutants and performed whole-genome sequencing and Sanger sequencing to investigate the potential EMB resistance mechanisms in *M. avium*.

## MATERIALS AND METHODS

### Strains and antibiotics

All the *M. avium* clinical strains were provided by The First Affiliated Hospital, Zhejiang University School of Medicine (Hangzhou, China). Clinical strains were identified as *M. avium* through whole-genome sequencing (Uni-medica, Shenzhen, China). Ethambutol (Aladdin, Shanghai, China) was dissolved in DMSO (Aladdin, Shanghai, China) and prepared as a 5.12 mg/mL stock solution and was stored at -20°C before use. Kanamycin sulfate (Aladdin, Shanghai, China) was dissolved in double distilled water and prepared as a 100 mg/mL stock solution, also was stored at -20°C before use.

### MIC determination and mutant screening

The MIC determination was performed using microdilution method as previously described (22). Drug solution was diluted to 128 µg/mL in Middlebrook 7H9 broth (Difco, USA) supplemented with 5% OADC (oleic acid, albumin, glucose, and catalase) and 0.2% glycerol, and serial two-fold dilutions (100 μL per well) were performed in 96-well plates (NEST, Wuxi, China). The bacterial suspension was diluted to 0.5 MacFarland Standard in Middlebrook 7H9 broth as mentioned above at first, then further diluted 100-fold, and a 100μL aliquot of the suspension was added into each well with EMB, resulting in 11 drug concentrations ranging from 64 μg/mL to 0.0625 μg/mL. The final bacterial count per well was approximately 5 x 10J CFU/mL (ranging from 1 x 10J ∼1 x 10^6^ CFU/mL). Positive control wells that contained only bacteria were set in the last column of the 96-well plates, and negative control wells that contained only drug were set separately. The plates were incubated at 37°C for 2 weeks, and the results were read. The MIC test was performed in triplicates.

After MIC determination in 7H9 broth, the strain used for isolating EMB-resistant mutant strains was identified. Middlebrook 7H11 agar plates (Hopebio, Qingdao, China, and containing 10% OADC and 0.5% glycerol) with EMB concentrations ranging from 1/4 to 4 times MIC were prepared, along with an antibiotic-free 7H11 agar plate as a control. The experimental approach for determining MIC on agar plates was described previously (23). A 20μL of 1000-fold diluted log-phase bacterial suspension was added to the 7H11 agar plates to confirm the MIC was consistent in both liquid and solid media. After 2 weeks of incubation at 37°C, the MIC on agar plates was determined.

For mutant screening, based on the MIC results from the 7H11 plates, 7H11 plates containing EMB at 4, 8, 16, and 32 times MIC were prepared. A 100μL aliquot of the stationary-phase bacterial suspension was then spread on the 7H11 EMB plates and incubated for 2 weeks to determine the optimal concentration to screen the EMB-resistant strains, which was used in the follow-up mutant screening. Single colonies of resistant mutants were picked and streaked onto 7H11 plates with higher concentration of EMB, followed by incubation for another two weeks to make sure that the bacteria are resistant to EMB. The original sensitive strain was streaked onto the plate as a control.

### Whole-genome sequencing

Genomic DNA was isolated from 10 mL of bacterial culture, and a number of resistant mutants and the original sensitive strain were picked. The cultures were centrifuged at 4000 rmp for 10 minutes at 4J°C, after which the supernatant was discarded. The pellets were washed twice with sterile water. The samples were sent for whole-genome sequencing (Novogene, Beijing, China). The genomic sequences of the resistant mutants were compared to those of the sensitive parent strain, and single nucleotide polymorphisms (SNPs) were analyzed using software Snippy to identify putative mutations associated with EMB resistance in *M. avium*.

### PCR and Sanger sequencing

The *ubiA* gene sequence was obtained from the whole-genome sequencing data of the susceptible strain by bacterial genome annotation software Bakta. Primers were designed as follows: ubiA_F: ATGACCGAGGAAACGCAGG; ubiA_R: CTAACCGAAGGCAACGGCG. Phanta Flash Master Mix (Dye Plus) (Vazyme, Nanjing, China) was used for PCR amplification, and the PCR conditions were as follows: denaturation at 95°C for 10 minutes, followed by 35 cycles of 95°C for 1 minute, 60°C for 1 minute, and 72°C for 30 seconds, with a final extension at 72°C for 1 minute.

For all remaining resistant mutants, the original susceptible strain and all other clinical strains, a 918 base pair fragment containing the *ubiA* gene was subjected to Sanger sequencing (Tsingke, Beijing, China), and the sequences were aligned for comparison by SnapGene.

### Gene complementation experiment

The target wild-type *ubiA* gene was amplified by PCR using the following primers: pmvubiA-F: caggaattcgatatcaagcttATGACCGAGGAAACGCAGG; pmvubiA-R: acgctagttaactacgtcgacCTAACCGAAGGCAACGGCG. The PCR product was purified by Fastpure Gel DNA Extraction Mini Kit (Vazyme). The pMV306hsp plasmid was digested with *Hind*III and *Sal*I, followed by cloning of the *ubiA* PCR product into the linearized plasmid vector using ClonExpress Ultra One Step Cloning Kit (Vazyme). The recombinant plasmid was then transformed into *E. coli* DH5α (Vazyme) by heat shock, and the transformant was incubated in a shaker at 37°C for 1 h followed by plating onto LB agar plates (Hopebio, Qingdao, China) containing 100 µg/mL kanamycin and incubated at 37°C overnight. Single colonies were selected the next day, cultured in LB broth (Hopebio, Qingdao, China) supplemented with 100 µg/mL kanamycin, and verified by Sanger sequencing to confirm that the wild-type *ubiA* gene was successfully cloned into pMV306hsp. The wild-type *ubiA* recombinant plasmid was then extracted with the Fastpure Plasmid Mini Kit (Vazyme) and electroporated (2,500 V, 1,000 Ω, and 25uF) into the EMB-resistant strains. The electroporated mutant strains were incubated in a 37°C shaker overnight and then plated onto 7H11 agar plates containing 100 µg/mL kanamycin and incubated at 37°C for 2 weeks. Single colonies that grew on the plates were then subjected to PCR sequencing to confirm that the wild-type *ubiA* gene was successfully introduced into the EMB-resistant strains. The complementation strains were tested for EMB susceptibility in 7H9 broth to validate the association between *ubiA* mutation and EMB resistance. Empty vector controls were included to exclude the potential effects of vector on MIC determinations.

## RESULTS

### Determination of EMB MICs for *M. avium* clinical strains

The EMB MIC values for 40 *M. avium* clinical strains were determined as described in the Methods section. As summarized in Table 1, according to the previous study (12), only 1 clinical strain (strain 245) was susceptible to EMB (MIC≤2 μg/mL), while all other clinical strains exhibited at least intermediate susceptibility (MIC≥4 μg/mL) to EMB (97.5%, 39/40), and 82.5% of the clinical strains (33/40) were resistant (MIC≥8 μg/mL) to EMB (12).

**Table 1.**
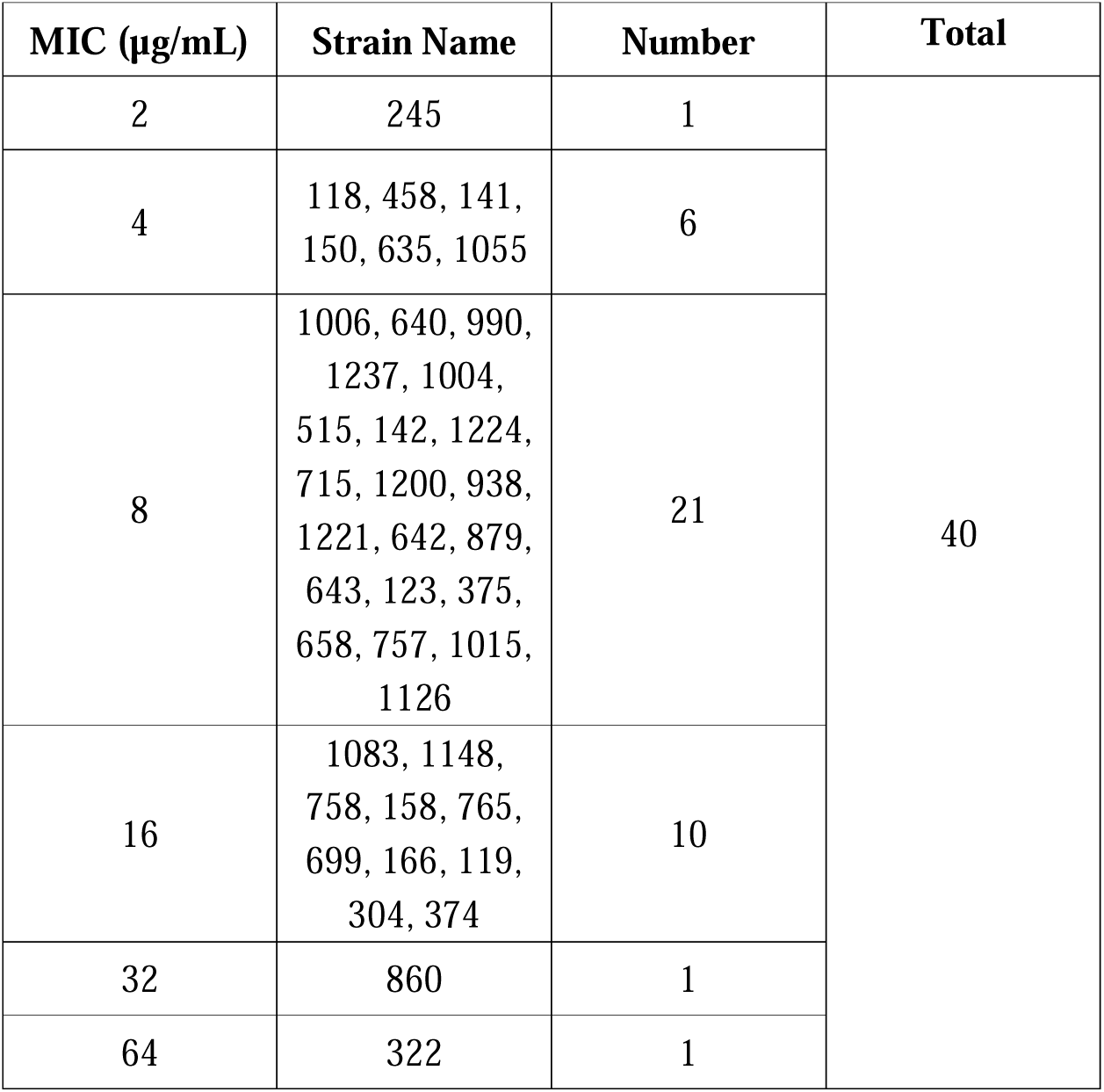
Ethambutol MIC distribution of *M. avium* clinical isolates.

### Mutant isolation

From the above results, we selected the EMB susceptible strain 245 (EMB MIC=2 μg/mL) for isolating EMB-resistant mutants. Since the isolation of mutant strains was performed on 7H11 plates, the MIC of clinical strain 245 was reassessed on 7H11 agar plates containing EMB at concentrations ranging from 0.5 to 8 μg/mL, based on the previous MIC result obtained in 7H9 broth. The final EMB MIC of strain 245 on 7H11 agar plate was determined to be 4 μg/mL, which is two-fold higher than its MIC in 7H9 liquid medium.

Based on the MIC results obtained from 7H11 agar plates, mutant isolation was conducted on 7H11 agar plates containing EMB at concentrations of 4X MIC and above, i.e., 16, 32, 64, and 128 μg/mL After 2 weeks of incubation, it was determined that 16 μg/mL was the optimal concentration for mutant isolation, as 7H11 agar plates containing higher EMB concentrations failed to grow mutant colonies. The concentration 16 μg/mL was subsequently used for large-scale screening of EMB-resistant mutants. All the colonies that grew on the plates containing 16 μg/mL were picked and streaked onto 7H11 agar plates containing EMB at 32 μg/mL together with the original susceptible strain. The mutants that grew both on plates containing 16 μg/mL and 32 μg/mL EMB were regarded as authentic resistant mutants, and finally a total of 121 mutants were successfully isolated.

### Whole-genome sequencing and Sanger sequencing

To identify the potential mechanisms of EMB resistance in *M. avium*, whole-genome sequencing was performed initially on eight EMB-resistant mutants along with the parent susceptible strain as a control. The results showed that all the mutants harbored mutations in the *ubiA* gene, strongly implying an association between *ubiA* gene mutation and EMB resistance in *M. avium*. The remaining 113 resistant mutants were all subsequently subjected to *ubiA* Sanger sequencing. The results showed that most of the mutants harbored mutations in *ubiA* (93.81%, 106/113), which is consistent with and supports the initial whole-genome sequencing findings. Integrating the results from both whole-genome and Sanger sequencing, the mutations were found to be distributed throughout the *ubiA* gene. The majority of mutants carried single-site point mutations at different locations of the *ubiA* gene, accounting for 80.99% (98/121) of the observed mutations. However, 13.22% (16/121) of the mutants had multi-site mutations, insertions, and deletions in *ubiA* gene (Table 2). Additionally, 7 mutants (5.79%, 7/121) exhibited no mutations in the *ubiA* gene. The results of the *ubiA* mutations identified through both whole-genome sequencing and Sanger sequencing are summarized in Table 2.

**Table 2.**
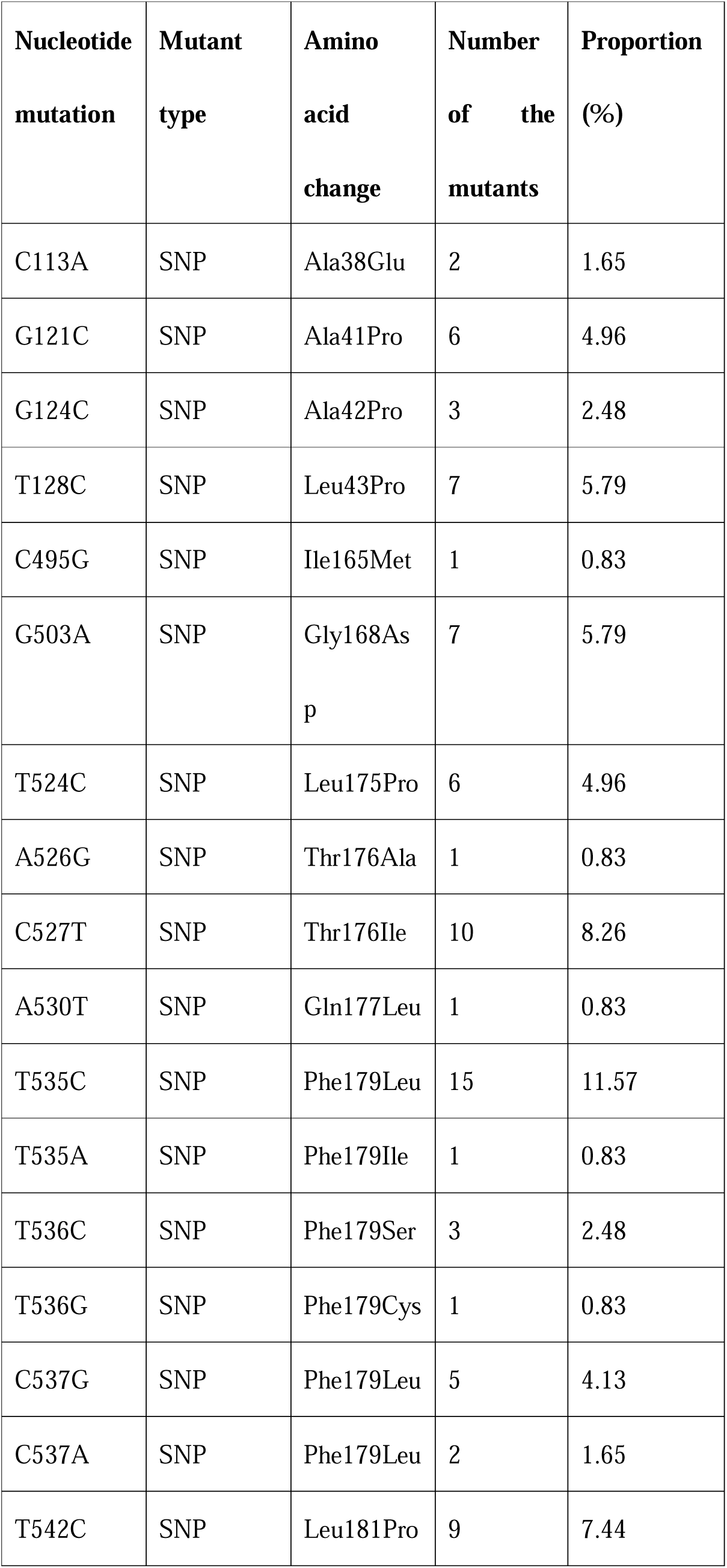
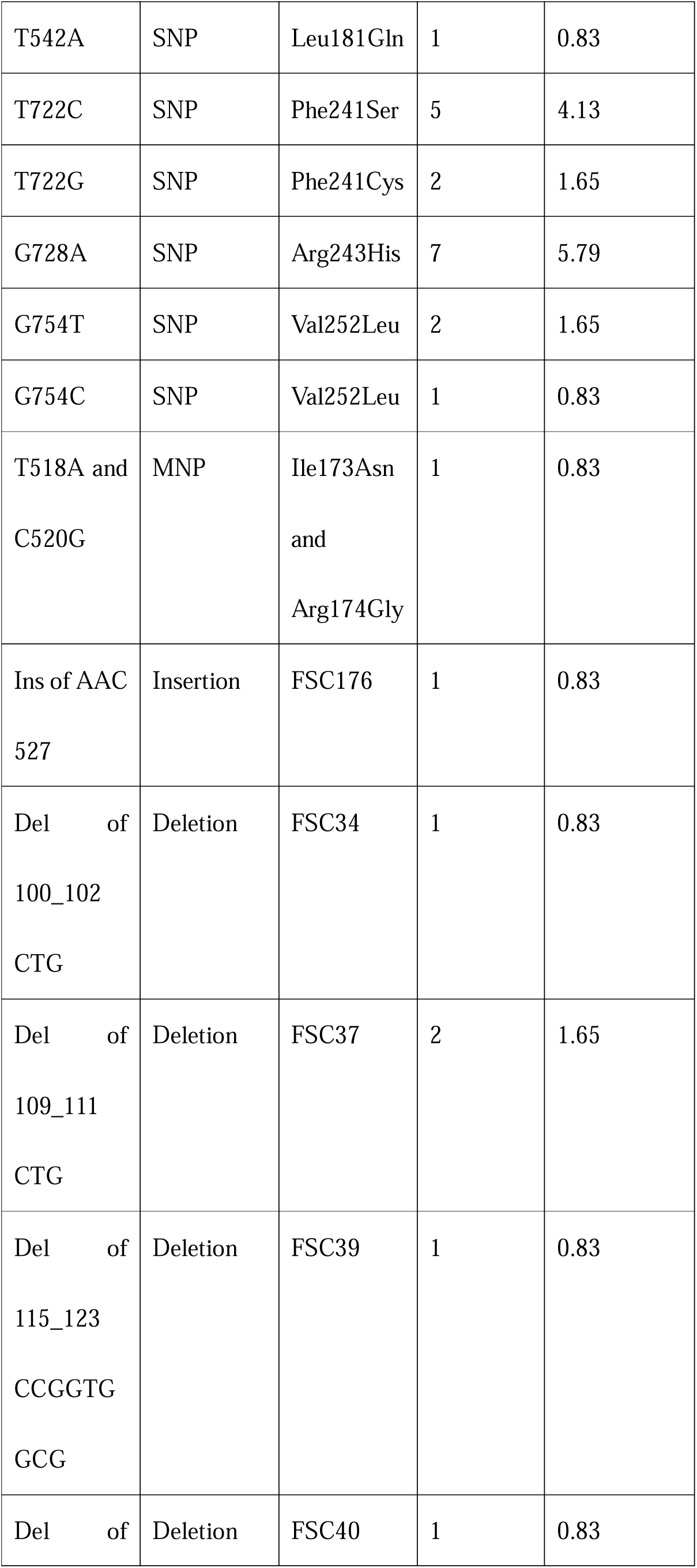
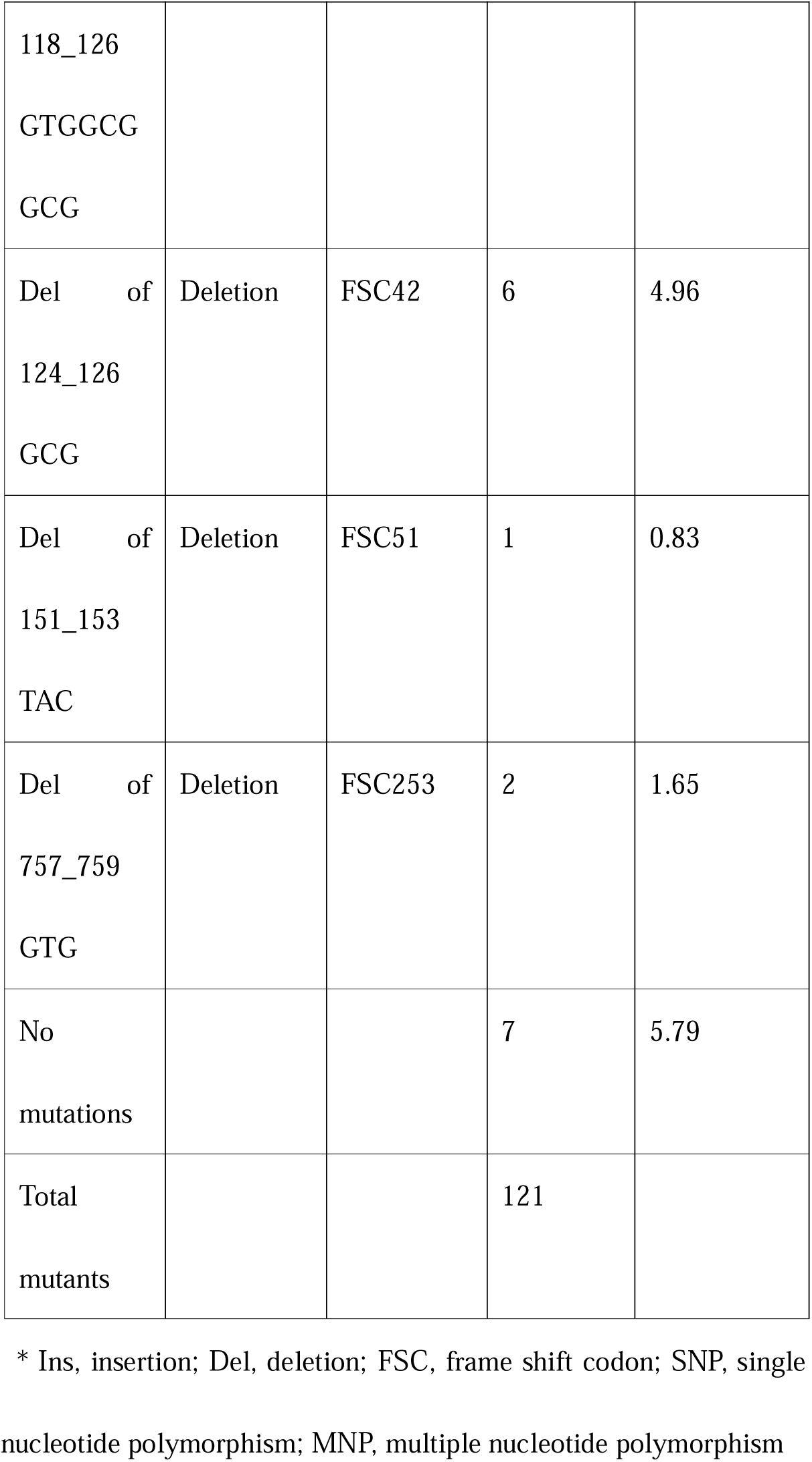
The proportion of *ubiA* mutations at different sites in all EMB-resistant mutants*.

### Majority of *M. avium* clinical isolates resistant to EMB carry mutations in *ubiA* gene

To assess whether mutations in *ubiA* were associated with EMB resistance in *M. avium* clinical strains, we performed Sanger sequencing of the *ubiA* gene on the remaining 39 clinical *M. avium* strains (Table 1) and compared their sequences with that of strain 245, the EMB-susceptible strain. Sequencing results revealed nonsynonymous *ubiA* mutations in 32 of 39 (82.05%) clinical strains, while 7 strains (6 strains with EMB MIC≥8 μg/mL, 1 strain with MIC=4 μg/mL) (7 of 39=17.95%) did not have *ubiA* mutations. Among the 32 strains with *ubiA* mutations, 27 were resistant to EMB (MIC≥8 μg/mL) and 5 exhibited intermediate susceptibility (MIC=4 μg/mL), suggesting a correlation between *ubiA* variations and EMB susceptibilities in different *M. avium* clinical strains. The Sanger sequencing results for the *ubiA* gene in clinical strains are summarized in Table 3.

**Table 3.**
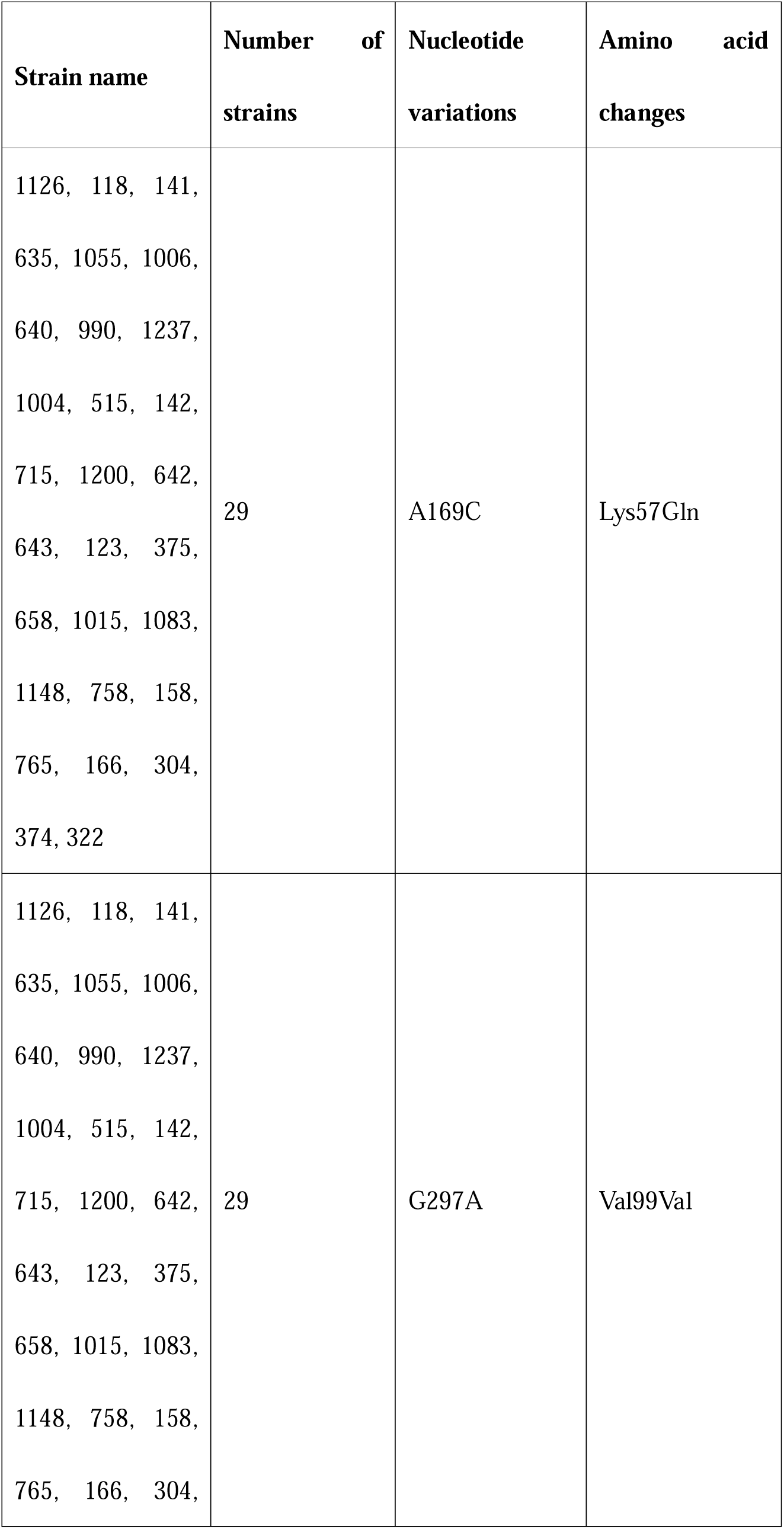
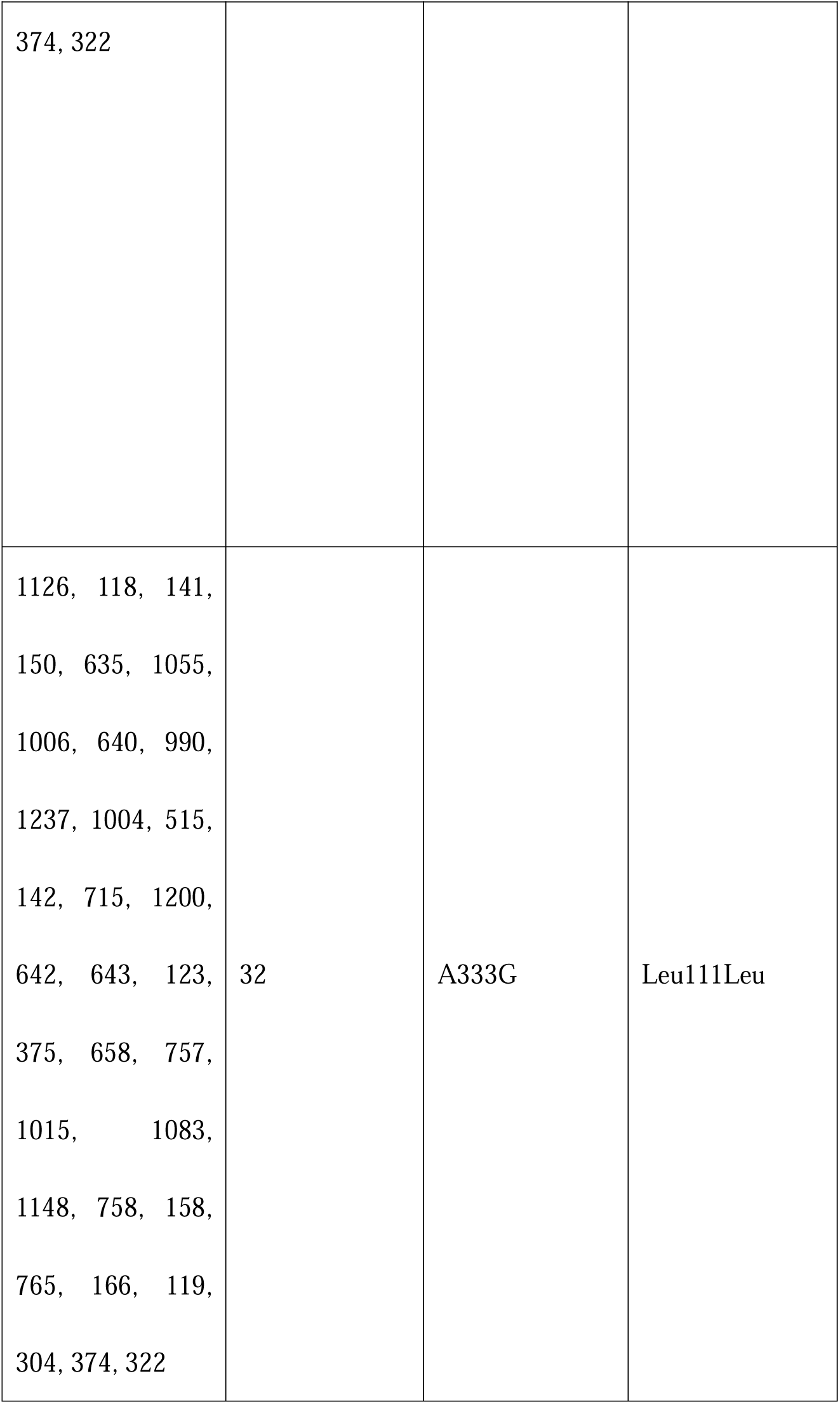
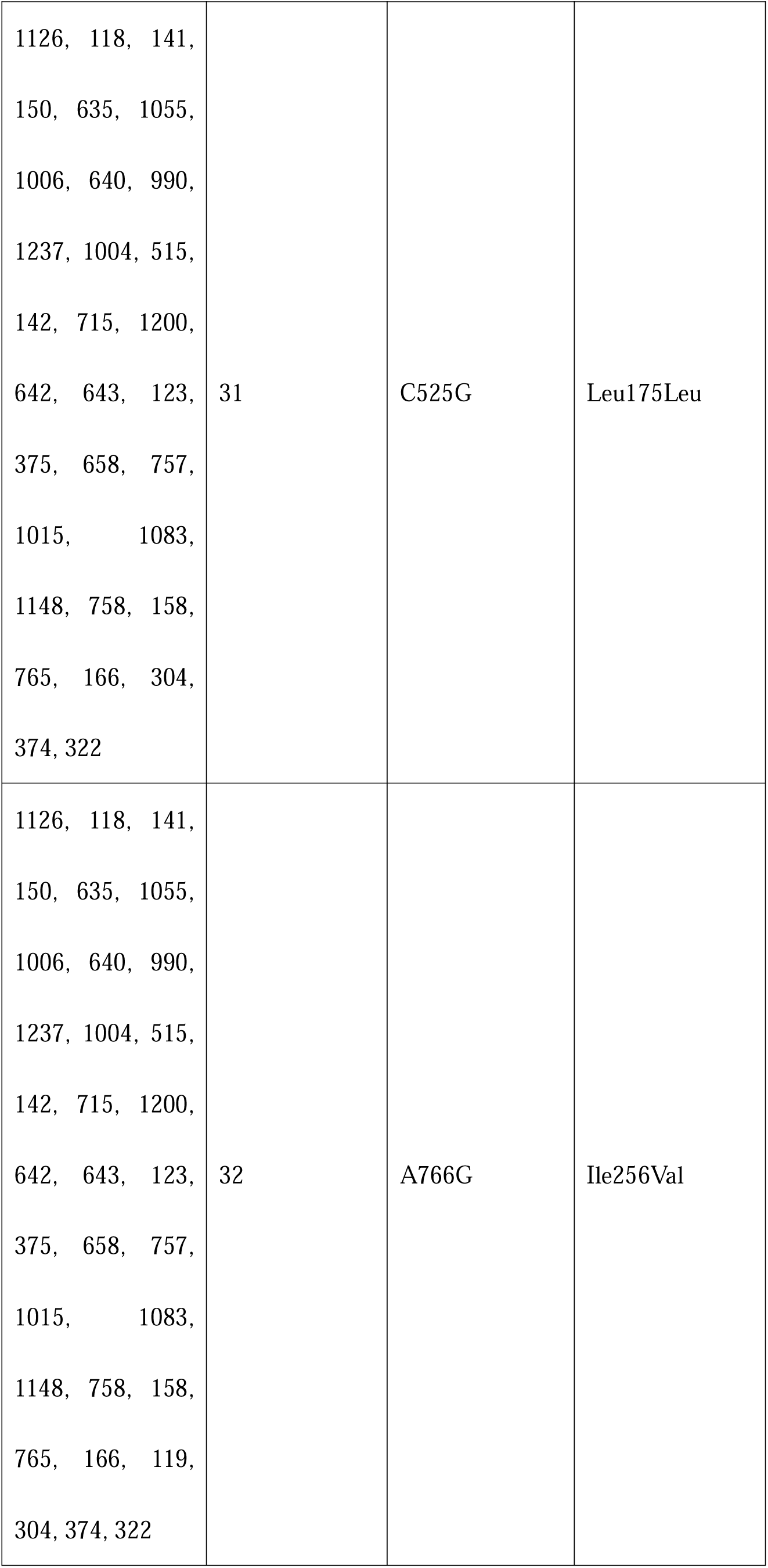
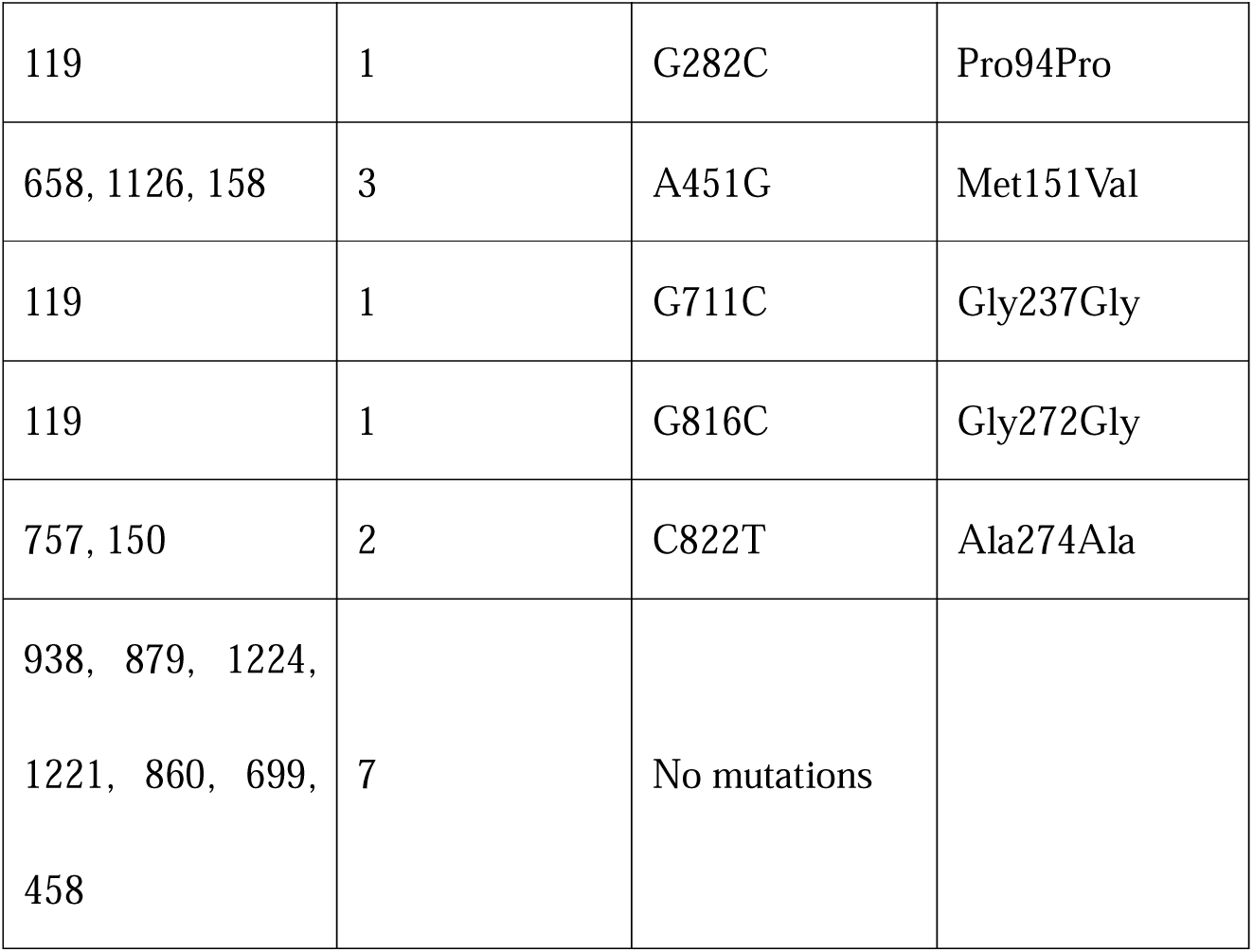
Mutations in the *ubiA* gene of *M. avium* clinical strains compared to the EMB susceptible strain 245.

### Complementation of the EMB-resistant mutant with the wild type *ubiA* restored EMB susceptibility

To determine whether the identified mutations in the *ubiA* gene contribute to EMB resistance, we subsequently introduced the wild-type *ubiA* gene into the randomly selected EMB-resistant mutants. Mutant strain 1 (Mutation: G124C) and Mutant strain 19 (Mutation: G754C) were used in the gene complementation experiment, with the empty vector being included as a control. The complemented mutants exhibited significantly reduced EMB MIC values (4 μg/mL) compared to mutants transformed with empty vector control (32 μg/mL) (Table 4). To determine whether the role of *ubiA* in EMB resistance is consistent across different genetic backgrounds, we further electroporated the wild-type *ubiA* gene from strain 245 into strain 322, which exhibited the highest EMB MIC (64 μg/mL) among the clinical strains tested in our study (Table 1). Complementation with the wild-type *ubiA* gene also restored EMB susceptibility, reducing the EMB MIC in strain 322 from 64 μg/mL to 8 μg/mL (Table 4). These results indicated that mutations in *ubiA* gene are directly responsible for EMB resistance in diverse *M. avium* strains.

**Table 4.**
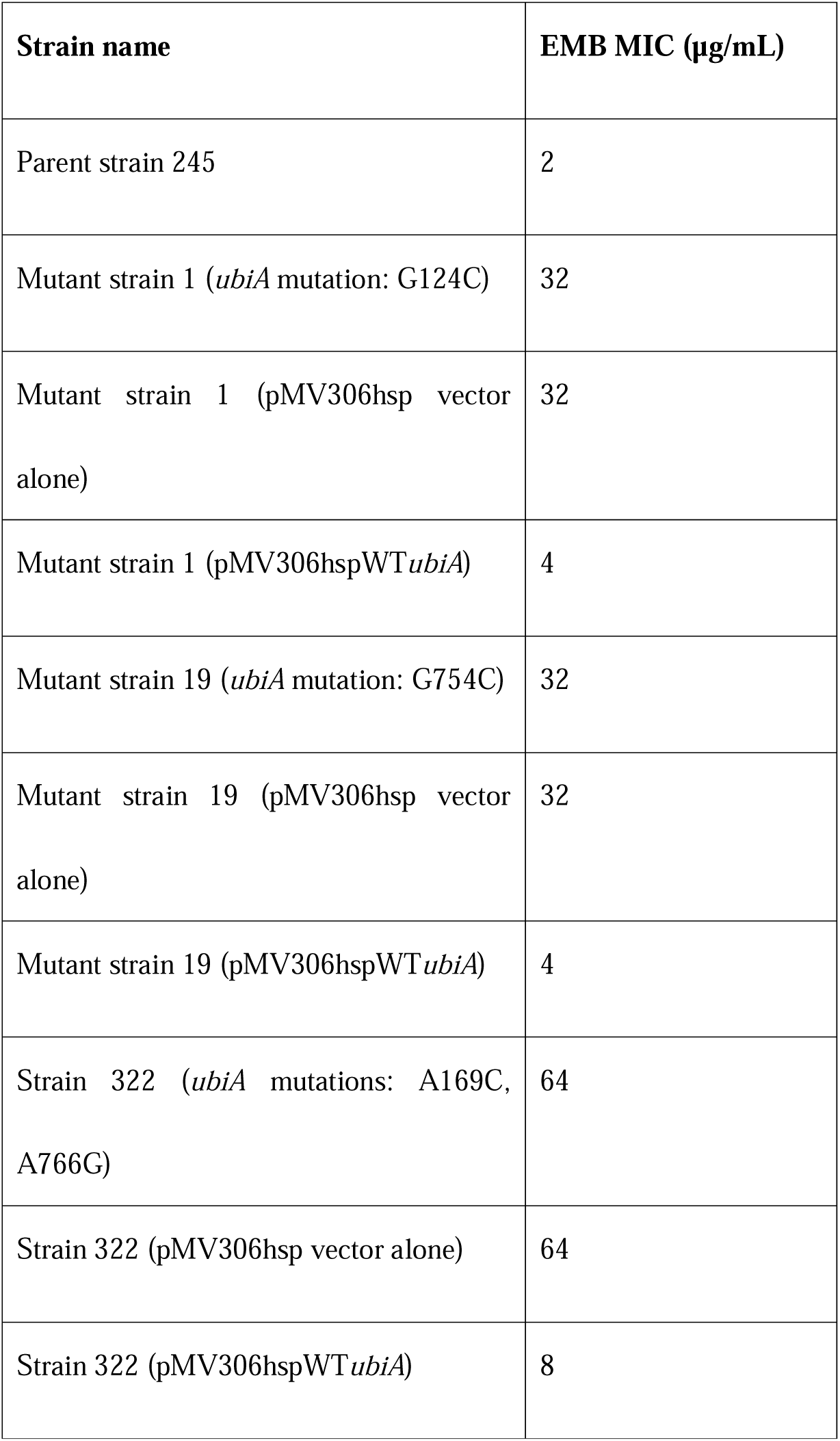
MICs of *M. avium* parent strain and *ubiA* mutant strains complemented with the wild-type *ubiA* gene.

## DISCUSSION

Although the use of EMB in the treatment of MAC infections is now widely accepted, the mechanisms underlying EMB resistance in this pathogen remain poorly characterized (8). In this study, we screened 121 EMB-resistant mutants of *M. avium* and performed whole-genome sequencing and targeted *ubiA* Sanger sequencing to investigate the genetic basis of resistance of *M. avium* to EMB. Integrated analysis of both whole-genome sequencing and *ubiA* sequencing data revealed that 94.21% (114/121) of the EMB-resistant strains harbored mutations in the *ubiA* gene, indicating that mutations in *ubiA* are the major mechanism of EMB resistance in *M. avium*. Furthermore, genetic complementation experiments confirmed that *ubiA* is the causative of EMB resistance in *M. avium*.

*ubiA*, which encodes DPPR (decaprenylphosphoryl-β-D-5-phosphoribose) synthase and corresponds to *Rv3806c* in *M. tuberculosis*, is involved in the synthesis of DPA (decaprenylphosphoryl-β-D-arabinose). DPA serves as the donor substrate for arabinosyltransferases including EmbB and EmbC, which are involved in catalyzing the transfer of arabinose from DPA to form the arabinan components of the mycobacterial cell wall (18, 24). Previous studies indicate that in strains with a wild-type *ubiA* genotype, EMB inhibits EmbCAB activity, thereby disrupting cell wall integrity and results in bacterial death (25, 26). However, when *ubiA* mutates, *ubiA* mutations will increase DPA levels, and the excess DPA will competitively binds to EmbCAB protein with EMB, reducing EMB inhibition and conferring high-level resistance in MTB (19). Given the close phylogenetic relationship between *M. avium* and MTB, it is plausible that *ubiA* mutations modulate EMB susceptibility through similar mechanisms in both species. However, in MTB, *ubiA* mutations actually account for a relatively small proportion of EMB-resistant strains and *embB* mutations are far more common (27). Even in high-level EMB-resistant MTB strains only, the proportion of *ubiA* mutation occurrence fluctuates with changes in geographical location but mostly does not exceed 10%, especially in Asian countries (28–31). Unlike MTB, *ubiA* mutations were identified in more than 90% of EMB-resistant *M. avium* mutants in this study.

It is noteworthy that 7 EMB-resistant mutants in this study lacked mutations in the *ubiA* gene. Furthermore, Sanger sequencing of *ubiA* gene in all 39 clinical strains revealed that 7 clinical strains, 6 of which exhibited EMB MICs≥8 μg/mL and one exhibited 4 μg/mL (Table 1 & 3), had no *ubiA* mutations compared with the susceptible control strain 245 (MIC=2 μg/mL). These findings strongly suggest the existence of alternative, as yet unidentified resistance mechanisms in *M. avium*, which warrants further investigations.

It is also worth noting that in our MIC assays, nearly all clinical strains exhibited resistance (MIC ≥ 8 μg/mL) or intermediate susceptibility (MIC=4 μg/mL) to EMB, based on breakpoints established in earlier study (12). Our MIC test results are consistent with the previous literature, enhancing the reliability of our results (32). This finding mentioned above stands in sharp contrast to the resistance profile observed in *M. tuberculosis*, where only 5.7% of newly diagnosed patients and 17.2% of re-treated patients show EMB resistance under the CLSI guidelines, which define susceptibility as MIC≤2 μg/mL and resistance as MIC≥8 μg/mL (33–35). The discrepancy of the EMB resistance rate between MTB and *M. avium* suggests that wild-type *M. avium* may commonly possess intrinsic resistance mechanisms against EMB, or the EMB’s susceptibility/resistance breakpoint for *M. avium* should be modified. Moreover, in contrast to EMB resistance in MTB, which is predominantly associated with mutations in *embB* (36), our results demonstrate that *ubiA* is the main contributor to EMB resistance in *M. avium*.

This study has several limitations. Firstly, as the *ubiA* gene is primarily associated with high-level EMB resistance, and all mutants in this study were selected on 7H11 agar plates containing 32 µg/mL EMB, our approach may have limited the detection of mutations conferring low-level EMB resistance. Secondly, most of the work in our study was conducted using a single clinical strain, and future studies should include a broader collection of EMB-resistant clinical isolates to further examine the correlation between *ubiA* mutations and EMB resistance in *M. avium*. Thirdly, We did not further investigate the resistance mechanisms of EMB-resistant clinical strains without *ubiA* gene mutations due to their different genetic backgrounds, making it difficult to identify the resistance mutations.

In conclusion, our study revealed that unlike MTB, where *embB* is the main resistance determinant*, ubiA* mutations are the major mechanism of EMB resistance in *M. avium*, which has not been reported before. The finding that *ubiA* mutations are the primary mechanisms of EMB resistance in *M. avium* has implications for rapid detection of EMB-resistant *M. avium* strains in the future. Further studies are still needed to investigate the alternative mechanisms of EMB resistance in *M. avium*.

## ACKNOWLEDGMENTS

This study was supported by National Infectious Disease Medical Center startup fund (YZ)(B2022011-1).

## Declaration

The authors declare that there is no conflict of interest regarding the publication of this article.

## Ethical approval

Not required.

## Data availability

Not applicable.

