## Supplementary material for "Mutations in the *ubiA* gene are the major mechanism of ethambutol resistance in *Mycobacterium avium*": Table 3

**Table 3. Mutations in the *ubiA* gene of *M. avium* clinical strains compared to the EMB susceptible strain 245**

| **Strain name** | **Number of strains** | **Nucleotide variations** | **Amino acid changes** |
| --- | --- | --- | --- |
| 1126, 118, 141, 635, 1055, 1006, 640, 990, 1237, 1004, 515, 142, 715, 1200, 642, 643, 123, 375, 658, 1015, 1083, 1148, 758, 158, 765, 166, 304, 374, 322 | 29 | A169C | Lys57Gln |
| 1126, 118, 141, 635, 1055, 1006, 640, 990, 1237, 1004, 515, 142, 715, 1200, 642, 643, 123, 375, 658, 1015, 1083, 1148, 758, 158, 765, 166, 304, 374, 322 | 29 | G297A | Val99Val |
| 1126, 118, 141, 150, 635, 1055, 1006, 640, 990, 1237, 1004, 515, 142, 715, 1200, 642, 643, 123, 375, 658, 757, 1015, 1083, 1148, 758, 158, 765, 166, 119, 304, 374, 322 | 32 | A333G | Leu111Leu |
| 1126, 118, 141, 150, 635, 1055, 1006, 640, 990, 1237, 1004, 515, 142, 715, 1200, 642, 643, 123, 375, 658, 757, 1015, 1083, 1148, 758, 158, 765, 166, 304, 374, 322 | 31 | C525G | Leu175Leu |
| 1126, 118, 141, 150, 635, 1055, 1006, 640, 990, 1237, 1004, 515, 142, 715, 1200, 642, 643, 123, 375, 658, 757, 1015, 1083, 1148, 758, 158, 765, 166, 119, 304, 374, 322 | 32 | A766G | Ile256Val |
| 119 | 1 | G282C | Pro94Pro |
| 658, 1126, 158 | 3 | A451G | Met151Val |
| 119 | 1 | G711C | Gly237Gly |
| 119 | 1 | G816C | Gly272Gly |
| 757, 150 | 2 | C822T | Ala274Ala |
| 938, 879, 1224, 1221, 860, 699, 458 | 7 | No mutations |  |
