## Supplementary material for "Mutations in the *ubiA* gene are the major mechanism of ethambutol resistance in *Mycobacterium avium*": Table 4

**Table 4. MICs of *M. avium* parent strain and *ubiA* mutant strains complemented with the wild-type *ubiA* gene**

| **Strain name** | **EMB MIC (µg/mL)** |
| --- | --- |
| Parent strain 245 | 2 |
| Mutant strain 1 (*ubiA* mutation: G124C) | 32 |
| Mutant strain 1 (pMV306hsp vector alone) | 32 |
| Mutant strain 1 (pMV306hspWT*ubiA*) | 4 |
| Mutant strain 19 (*ubiA* mutation: G754C) | 32 |
| Mutant strain 19 (pMV306hsp vector alone) | 32 |
| Mutant strain 19 (pMV306hspWT*ubiA*) | 4 |
| Strain 322 (*ubiA* mutations: A169C, A766G) | 64 |
| Strain 322 (pMV306hsp vector alone) | 64 |
| Strain 322 (pMV306hspWT*ubiA*) | 8 |
