## Supplementary material for "Mutations in the *ubiA* gene are the major mechanism of ethambutol resistance in *Mycobacterium avium*": Table 1

**Table 1. Ethambutol MIC distribution of *M. avium* clinical isolates**

| **MIC (µg/mL)** | **Strain Name** | **Number** | **Total** |
| --- | --- | --- | --- |
| **2** | 245 | 1 | 40 |
| **4** | 118, 458, 141, 150, 635, 1055 | 6 |  |
| **8** | 1006, 640, 990, 1237, 1004, 515, 142, 1224, 715, 1200, 938, 1221, 642, 879, 643, 123, 375, 658, 757, 1015, 1126 | 21 |  |
| **16** | 1083, 1148, 758, 158, 765, 699, 166, 119, 304, 374 | 10 |  |
| **32** | 860 | 1 |  |
| **64** | 322 | 1 |  |
