## Supplementary material for "Mutations in the *ubiA* gene are the major mechanism of ethambutol resistance in *Mycobacterium avium*": Table 2

**Table 2.** **The proportion of *ubiA* mutations at different sites in all EMB-resistant mutants***

| **Nucleotide mutation** | **Mutant type** | **Amino acid change** | **Number of the mutants** | **Proportion (%)** |
| --- | --- | --- | --- | --- |
| C113A | SNP | Ala38Glu | 2 | 1.65 |
| G121C | SNP | Ala41Pro | 6 | 4.96 |
| G124C | SNP | Ala42Pro | 3 | 2.48 |
| T128C | SNP | Leu43Pro | 7 | 5.79 |
| C495G | SNP | Ile165Met | 1 | 0.83 |
| G503A | SNP | Gly168Asp | 7 | 5.79 |
| T524C | SNP | Leu175Pro | 6 | 4.96 |
| A526G | SNP | Thr176Ala | 1 | 0.83 |
| C527T | SNP | Thr176Ile | 10 | 8.26 |
| A530T | SNP | Gln177Leu | 1 | 0.83 |
| T535C | SNP | Phe179Leu | 15 | 11.57 |
| T535A | SNP | Phe179Ile | 1 | 0.83 |
| T536C | SNP | Phe179Ser | 3 | 2.48 |
| T536G | SNP | Phe179Cys | 1 | 0.83 |
| C537G | SNP | Phe179Leu | 5 | 4.13 |
| C537A | SNP | Phe179Leu | 2 | 1.65 |
| T542C | SNP | Leu181Pro | 9 | 7.44 |
| T542A | SNP | Leu181Gln | 1 | 0.83 |
| T722C | SNP | Phe241Ser | 5 | 4.13 |
| T722G | SNP | Phe241Cys | 2 | 1.65 |
| G728A | SNP | Arg243His | 7 | 5.79 |
| G754T | SNP | Val252Leu | 2 | 1.65 |
| G754C | SNP | Val252Leu | 1 | 0.83 |
| T518A and C520G | MNP | Ile173Asn and Arg174Gly | 1 | 0.83 |
| Ins of AAC 527 | Insertion | FSC176 | 1 | 0.83 |
| Del of 100_102 CTG | Deletion | FSC34 | 1 | 0.83 |
| Del of 109_111 CTG | Deletion | FSC37 | 2 | 1.65 |
| Del of 115_123 CCGGTGGCG | Deletion | FSC39 | 1 | 0.83 |
| Del of 118_126 GTGGCGGCG | Deletion | FSC40 | 1 | 0.83 |
| Del of 124_126 GCG | Deletion | FSC42 | 6 | 4.96 |
| Del of 151_153 TAC | Deletion | FSC51 | 1 | 0.83 |
| Del of 757_759 GTG | Deletion | FSC253 | 2 | 1.65 |
| No mutations |  |  | 7 | 5.79 |
| Total mutants |  |  | 121 |  |

* Ins, insertion; Del, deletion; FSC, frame shift codon; SNP, single nucleotide polymorphism; MNP, multiple nucleotide polymorphism
